# Mutation of the *Drosophila* serotonin transporter dSERT disrupts courtship and feeding and increases both daytime and nighttime sleep

**DOI:** 10.1101/2022.06.10.495593

**Authors:** Elizabeth M. Knapp, Andrea Kaiser, Rebecca C. Arnold, Maureen M. Sampson, Manuela Ruppert, Li Xu, Matthew I. Anderson, Shivan L. Bonanno, Henrike Scholz, Jeffrey M. Donlea, David E. Krantz

## Abstract

The Serotonin Transporter (SERT) regulates extracellular serotonin levels and is the target of most current drugs used to treat depression. The mechanisms by which inhibition of SERT activity influences behavior are poorly understood. To address this question in the model organism *Drosophila melanogaster*, we developed new loss of function mutations in *Drosophila SERT* (*dSERT).* Previous studies in both flies and mammals have implicated serotonin as an important neuromodulator of sleep, and our newly generated *dSERT* mutants show an increase in total sleep and altered sleep architecture. Differences in daytime vs. nighttime sleep architecture as well as genetic rescue experiments unexpectedly suggest that distinct serotonergic circuits may modulate daytime versus nighttime sleep. *dSERT* mutants also show defects in copulation and food intake, akin to the clinical side effects of SSRIs. Starvation did not overcome the sleep drive in the mutants. Additionally in males, but not female *dSERT* mutants, the drive to mate also failed to overcome sleep drive. *dSERT* may be used to further explore the mechanisms by which serotonin regulates sleep and its interplay with other complex behaviors.

**Author Summary:** Many medications used to treat depression and anxiety act by changing serotonin levels in the brain. Fruit flies also use serotonin and can be used as a model to study the brain. We have made a fly mutant for the serotonin transporter (SERT), which is the target of antidepressants in humans. The mutants sleep more, eat less, and have a decreased sex drive. These flies can be used to study the neuronal pathways by which serotonin regulates sleep, eating and sexual behaviors and may help us to understand the behavioral effects of antidepressants.

## Introduction

Sleep is essential for life and is evolutionarily conserved from insects to mammals (1–3). Both the amount and architecture of sleep are critical for cognition and disruption of sleep in humans is linked to neurological and psychiatric disorders (4–6). The neuromodulator serotonin (5-hydroxytryptamine, 5-HT) acts as a key regulator of sleep, and its involvement in sleep has been known for decades (7, 8). Previous studies in vertebrates have shown that serotonin signaling can promote wakefulness, while paradoxically, others demonstrate that serotonin is critical for sleep induction and maintenance (9–13). The circuits responsible for these effects and the cellular mechanisms by which serotonin regulates sleep remain unclear.

For over two decades, *Drosophila* has been utilized as a model system to study sleep (14, 15) and both serotonin (16–20) and dopamine (21–25) have been shown to play significant roles in regulating sleep duration and quality. A role for serotonin in promoting sleep in *Drosophila* has been demonstrated in part by feeding the precursor 5-hydroxytryptophan (5-HTP) (16) or by utilizing mutants for the serotonin rate-limiting synthetic enzyme tryptophan hydroxylase (TRH) (20). Furthermore, of the five serotonin receptors expressed in *Drosophila*, mutations in three (*d5-HT1A*, *d5-HT2B* and *d5-HT7*) disrupt sleep behaviors, including total sleep amount, sleep rebound, and sleep architecture (16,18,20). The powerful molecular genetic tools available in the fly have allowed some of the specific structures and cells required for sleep to be identified (16,20,21,26–32). Similarly, these tools have the potential to uncover the mechanisms by which specific circuits regulate sleep.

The primary mechanism by which serotonin is cleared from the extracellular space in both mammals and flies is reuptake via the Serotonin Transporter (SERT) which localizes to the plasma membrane of serotonergic neurons (33). Blockade of SERT activity increases the amount of serotonin available for neurotransmission. It is the primary target for most current treatments of depression and anxiety disorders, including the widely prescribed Selective Serotonergic Reuptake Inhibitors (SSRIs) (34). In addition to their therapeutic effects, SSRIs can dramatically influence a variety of other physiological functions and behaviors such as eating, libido, and sleep (35–38). Consistent with the complex relationship between serotonin and sleep, SSRIs often produce diverse and sometimes contradictory defects on sleep including insomnia, decreased REM sleep efficiency and daytime somnolence (9,38,39). It remains unclear how changes in SERT activity modifies sleep behavior in mammals.

A relatively weak hypomorph of *dSERT* has been previously described (40). The *dSERT* MiMIC insertion lies within the first intron just upstream of the first coding exon of the gene and reduces *dSERT* mRNA expression by ∼50% (40). Its potential effects on sleep were not reported, and in general, it can be difficult to make firm conclusions about mutations that are not null or at least strong hypomorphs.

In this study, we have used P-element excision to generate new mutant alleles and find that *dSERT* is required for regulating both sleep amount and architecture. Our work further elucidates how the increased sleep drive exhibited in these *dSERT* mutants is affected in the context of other critical behaviors that are regulated by serotonin signaling including mating and feeding. Our data also suggest that *dSERT’s* role in daytime vs. nighttime sleep may be modulated via distinct serotonergic circuits.

## Results

### Disruption of *dSERT* increases sleep drive in *Drosophila*

To generate new mutant alleles of *dSERT*, we used imprecise excision of a transposable element in the line *XP^d04388^* (BDSC #85438). To ensure a consistent genetic background, *XP^d04388^* was first outcrossed into *w^1118^*, and *w^1118^* served as the primary control for our initial assays. The proximal end of the P element in *XP^d04388^* is 514 bp 5’ of the predicted transcriptional start site of *dSERT* (Figure 1A). We screened for loss of the *white* minigene in the P element and recovered two imprecise excision alleles of 1121bp and 1178bp which we designate as *dSERT^10^* and *dSERT^16^* respectively (Figure 1A). Both *dSERT^10^* and *dSERT^16^* delete the first non-coding exon and the first intron of *dSERT*. We identified two additional lines as controls; *dSERT^1^* contains 41 additional bases that are remnants of the former P-element insertion but does not otherwise disrupt the *dSERT* gene and *dSERT^4^* with no detectable genomic alterations at the former P-element insertion side.

**Figure 1.**
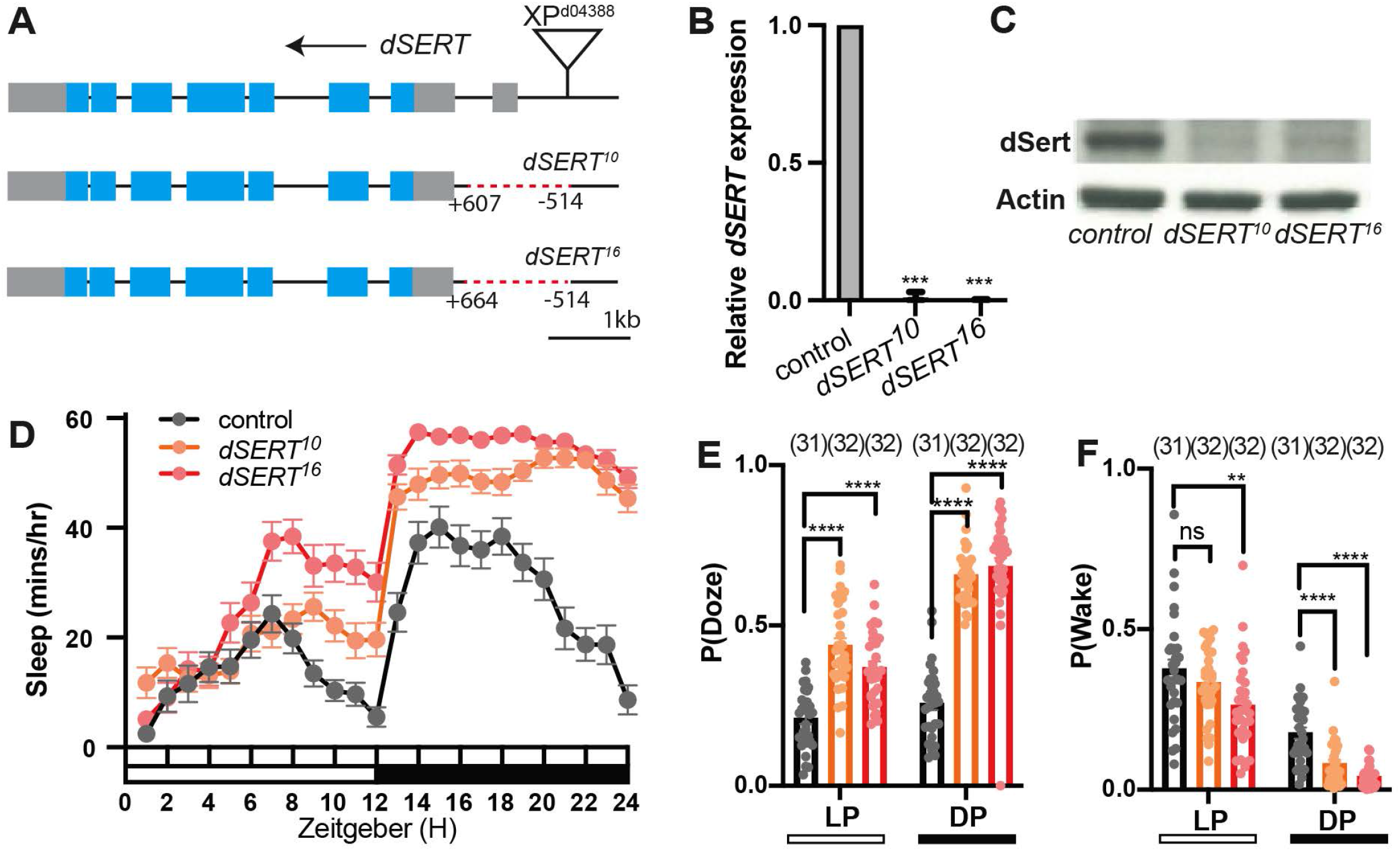
*dSERT* imprecise excision alleles. (A) Schematic of *dSERT* gene: grey and blue boxes indicate non-coding exons and coding exons respectively. Deletions of the *dSERT^10^* and *dSERT^16^* flies are indicated with red dotted lines. (B) qRT-PCR was performed on cDNA synthesized from whole male flies with *RplP0* for reference. Gene expression of *dSERT* in the mutant was normalized to controls with +1 representing expression in *w^1118^*. *dSERT* transcript levels were significantly down-regulated in *dSERT^10^* (0.01±0.02) and *dSERT^16^* (0.001±0.003). Error bars represent SD. P***≤0.001 (Student’s t-test). (C) Western blot analysis shows a band representing dSERT at ∼65kD that is reduced in intensity in both mutants. Actin was used as a loading control and shows no difference across genotypes. (D) Hourly sleep traces in wild-type *w^1118^* (black) *dSERT^10^* (orange) and *dSERT^16^* (red) flies. (E-F) Quantification of P(Doze) (E) and P(Wake) (F) during the light period (LP) and dark period (DP) in control *w^1118^*(grey), *dSERT^10^* mutants (orange), and *dSERT^16^*mutants (red). Sleep trace shows mean ± SEM and graphs show individual datapoints and group means ± SEM. P****≤0.0001 (one-way ANOVA with Holm-Sidak multiple comparisons tests).

To determine whether expression of *dSERT* was disrupted in *dSERT^10^* and *dSERT^16^*, we first used qRT-PCR to quantify mRNA expression, using *w^1118^* as a control. The *dSERT* transcript was not detectable in either of the mutant alleles (Figure 1B). By contrast, a previously published mutant allele generated by insertion of a MiMIC cassette was reported to retain ∼50% expression relative to wild type (40). To confirm these results and determine whether dSERT protein was similarly reduced, we performed western blots using a previously validated antibody to dSERT (41). Compared to *w^1118^* controls, both *dSERT^10^*and *dSERT^16^* show a decrease in dSERT protein expression (Figure 1C). It is unclear whether a faint band present in the western blots represents residual protein or non-specific background, and the intact coding sequence suggests that they may not be null alleles. Nonetheless, since they appeared to represent relatively strong hypomorphs with undetectable mRNA levels we preceded with our behavioral analyses.

We analyzed sleep in the *dSERT* mutants and found that both *dSERT^10^* and *dSERT^16^* mutants have drastically increased sleep compared to *w^1118^* controls (Figure 1D). The probability of transitioning from an awake to a sleep stake, P(Doze), and the probability of transition from a sleep to an awake state, P(Wake), were calculated to further analyze the changes in sleep drive and arousal in the *dSERT* mutants. Compared to controls, both *dSERT* mutants show a significant increase in P(Doze) (Figure 1E) and a reduction in P(Wake) (Figure 1F).

To confirm the increased sleep phenotypes are the result of *dSERT* disruption and not from other spurious changes to the genetic background, transheterozygous *dSERT* mutants (*dSERT^10^/dSERT^16^*) were assayed and shown to exhibit a significant increase in total sleep compared to controls (Sup Figure 1A-B). Consistent with these findings, *dSERT^16^* homozygotes also demonstrated a significant increase in total sleep when compared either to heterozygous *dSERT^16^*flies (Sup Figure 1C-D) or the revertant *dSERT^4^* control (Sup Figure 1E-F). Taken together, these results demonstrate disruption of *dSERT* causes a significant increase in overall sleep and sleep drive.

### Loss of *dSERT* significantly increases sleep and alters sleep architecture in daytime and nighttime

The sleep phenotype we observed was stronger in *dSERT^16^* compared to *dSERT^10^*, and we therefore focused on *dSERT^16^* for further analysis. To verify that the phenotype was not due to a bias in the time spent on one end of the testing tube or to limited movements that do not carry the flies past the tube’s midline, we assayed activity with both single-beam and multibeam monitors (42). Our results show that in both single-beam and multibeam monitors *dSERT^16^*mutants exhibit a significant increase in total sleep compared to controls (Figure 2A). These data eliminate the potential confound that can be caused by artefactual retention of the flies at one end of the assay tubes.

**Figure 2.**
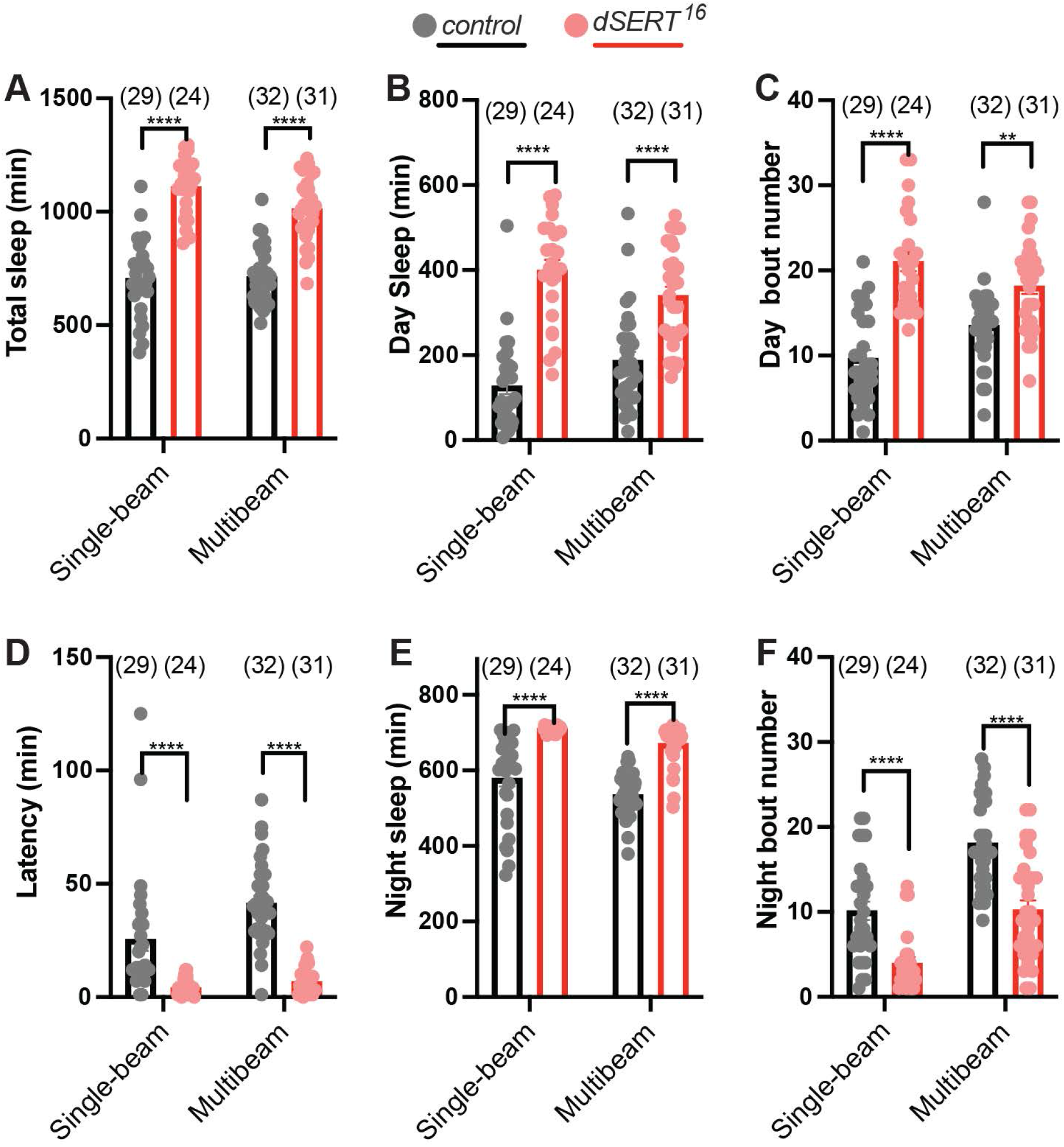
*dSERT^16^* mutants exhibit increased sleep behavior and changes in sleep architecture. Sleep was recorded with both single-beam and multibeam monitors. (A) Quantification of total sleep in wild-type *w^1118^* (grey) and *dSERT^16^* (red) flies. Analysis of daytime sleep (B) and daytime bout frequency (C). (D) Latency after light-off was significantly decreased in *dSERT^16^* mutants. Quantification of nighttime sleep (E) and nighttime bout frequency (F). Graphs show individual datapoints and group means ± SEM. Student’s t-test, two-way, unpaired, p≤0.0001****

Previous studies have shown that stimulation of aminergic pathways can increase grooming and alter other sensorimotor behaviors that might effect sleep (43–45). We therefore tested whether grooming or negative geotaxis were altered in the *dSERT* mutants. We did not detect any differences from controls in either grooming rate (Sup Figure 2A) or performance of negative geotaxis in *dSERT^16^*flies (Sup Figure 2B). Overall, these findings indicate that the behavioral change we detect represents an increased sleep in *dSERT* mutants rather than an artifact caused by changes in other amine-associated behaviors.

More specific analysis of both daytime and nighttime sleep behavior shows that *dSERT^16^* flies have dramatically increased sleep during the daytime (Figure 2B). This is further characterized by an increase in daytime bout frequency compared to controls (Figure 2C). In addition, *dSERT^16^*mutants also displayed reduced latency to sleep at night (Figure 2D). The analysis of nighttime sleep behavior revealed that *dSERT^16^* mutants similarly exhibit dramatically increased nighttime sleep (Figure 2E) but in contrast to their daytime behavior, we observed significant decrease in nighttime bout frequency (Figure 2F). Together, these findings indicate that disruption of *dSERT* causes a significant increase in sleep and alters sleep architecture in both the day and evening periods.

### *dSERT^16^* mutants exhibit rhythmic circadian behaviors

In addition to sleep, serotonergic pathways in mammals have been implicated in the regulation of circadian rhythmicity (46). To investigate a possible role for dSERT activity in sustaining circadian rhythms, we analyzed circadian locomotor behaviors in *dSERT^1^*^6^ mutants. Both control and *dSERT^16^* mutants exhibited robust locomotor rhythms with bimodal activity peaks in 12h/12h light/dark (LD) cycles (Figure 3A-B) and their rhythmic behaviors persisted in free-running constant darkness (DD) cycles (Figure 3E-F). Consistent with the increased sleep behavior of *dSERT^16^* mutants, the average activity of *dSERT^16^*flies was significantly decreased throughout both LD (Figure 3C) and DD (Figure 3G) conditions. Although the average activity was dampened, *dSERT^16^* mutants did not exhibit a significant decrease in either morning or evening anticipation behaviors for either LD (Figure 3D) or DD (Figure 3H) cycles. Periodogram analysis confirmed that *dSERT^16^*mutants behave indistinguishably from wildtype controls (Figure 3 I-J) and quantification of free-running circadian behaviors demonstrates that over 96% of *dSERT^16^* flies (tau=23.6 ±0.04) were rhythmic (Figure 3K) similar to control flies (tau=23.5±0.01). In addition, the circadian periods of *dSERT^16^* mutants did not appear significantly changed (Figure 3L). In summary, we find that loss of *dSER*T does not appear to disrupt circadian rhythmicity. These data are consistent with previous work showing the feeding of either 5-HTP, the SSRI fluoxetine (Prozac), or overexpression of *d5-HT1B* in flies also did not alter rhythms in free-running conditions (17).

**Figure 3.**
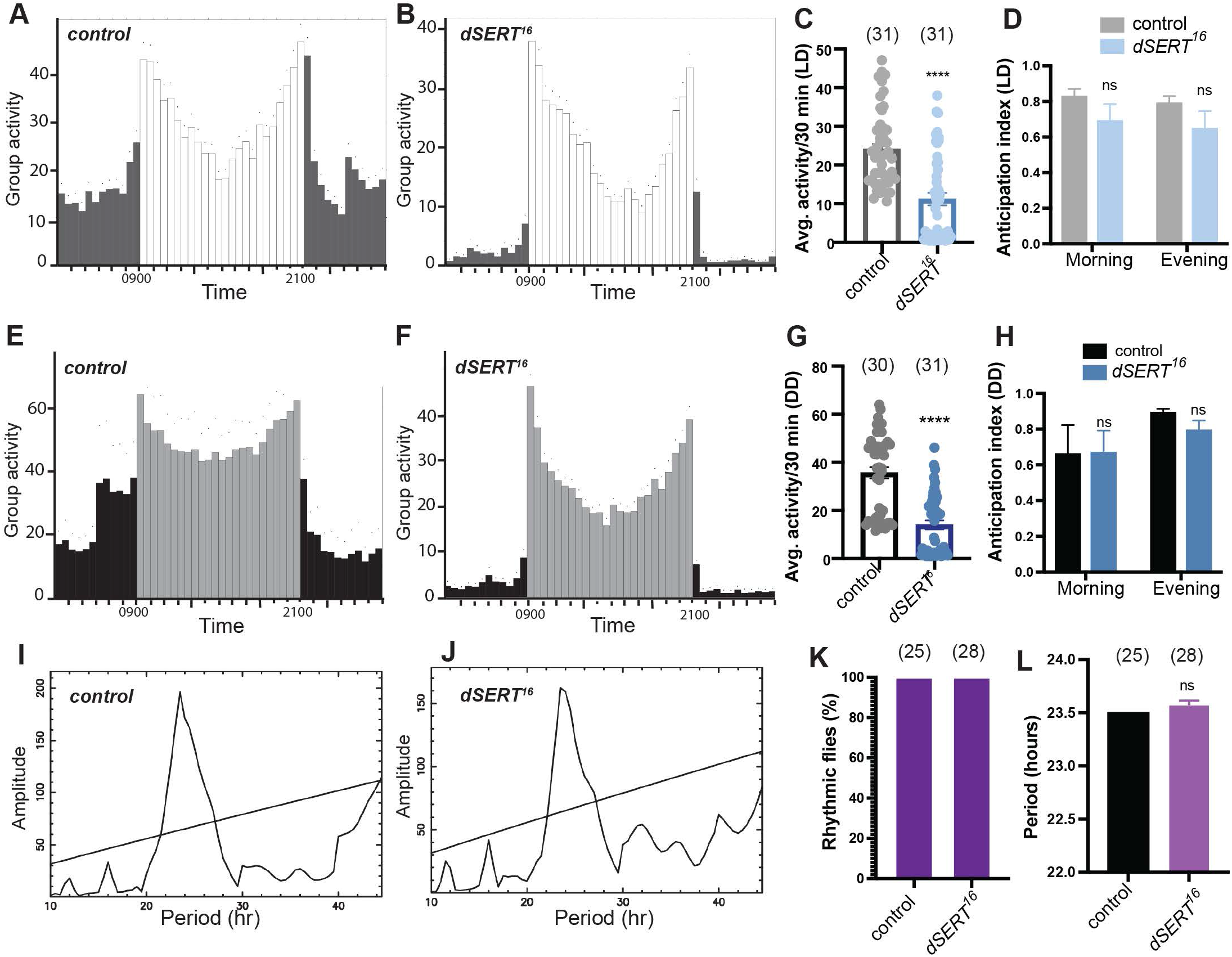
*dSERT^16^* mutants exhibit rhythmic circadian behaviors. (A-B) 12h:12h LD and (E-F) 12h:12h DD locomotor activity group analysis for *w^1118^* (A,E) and *dSERT^16^* (B,F) flies. Histograms represent the distribution of activity through 24 h, averaged over three days, number of flies indicated in C, D, G, H. Lighter and darker bars indicate day and night phases, respectively. Dots indicate the SEM of the activity for each 0.5 hr (0900 is lights-on and 2100 is lights-off). (C,G) Mean daily activity for *dSERT^16^* mutants is reduced during both LD (C) and DD (G) cycles, individual datapoints and group means ± SEM. Student’s t-test, two-way, unpaired, p****≤0.0001. Calculation of morning and evening anticipation indexes for *dSERT^16^* and control flies in LD (D) and DD (H) cycles. (I-J) Representative periodograms derived from activity records of individual *w^1118^* (I) and *dSERT^16^* (J) flies in constant darkness. (K) Percentage of flies with detectable rhythmicity was calculated for control and *dSERT^16^*. (L) Circadian periods in DD were averaged from rhythmic flies per each genotype, error bars indicate SEM.

### Loss of *dSERT* disrupts courtship and copulation behaviors

In clinical settings, SSRIs have number of side effects including sexual dysfunction, raising the possibility that loss of *dSERT* could potentially influence these activities in the fly. We first investigated whether *dSERT^16^* mutants show defects in courtship and copulation and how this would interact with sleep. Previous work demonstrated that pairing male and females flies together overnight dramatically suppresses sleep as a result of increased courtship activity (47–49). Consistent with these findings, co-housing pairs of wildtype male and female flies together resulted in a significant decrease in nighttime sleep compared to sleep recorded from isolated male or female flies (Figure 4A). However, pairing male and female *dSERT^16^* mutants together did not result in any significant sleep loss compared to isolated conditions (Figure 4A). These results indicate the increased sleep drive outweighs copulation drive in *dSERT^16^*mutants and suggest that loss of dSERT might decrease sexual activity.

**Figure 4.**
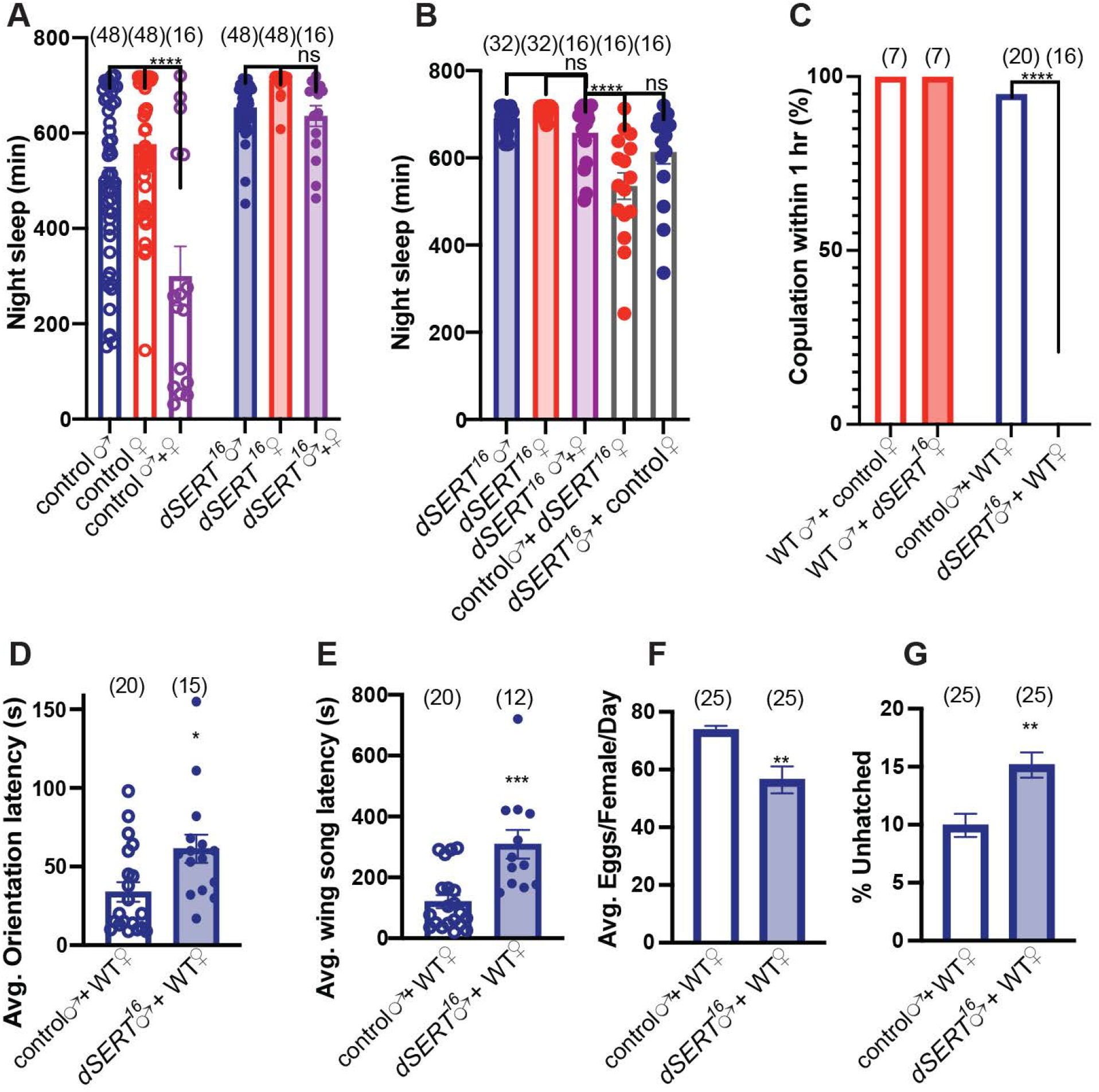
*dSERT^16^* mutants exhibit defects in courtship and copulation. (A) Quantification of nighttime sleep for males (blue) and females (red) individually housed in DAMs tubes for both control *w^1118^* (open bars/circles) and *dSERT^16^* (shaded bars/circles). Following 2 days of individual housing male and females were paired together in DAMs tubes (purple) and nighttime sleep was quantified for both *w^1118^*(open bar/circles) and *dSERT^16^* (shaded bar/circles). (B) Quantification of nighttime sleep for *dSERT^16^* males (shaded blue) and females (shaded red) individually housed or in male-female pairs (shaded purple). Nighttime sleep averages are also shown for mixed co-housing of control males with *dSERT^16^* females (open bars with red circles) or *dSERT^16^* males with control females (open bars with blue circles). (C-G) Mating pairs made up of either wildtype (WT) males (red) or WT females (blue) with controls (open bars) or *dSERT^16^* mutants (shaded bars). (C) Percentage of pairs that copulated within 1 hr. Quantification of average latency to orientation (D), and wing song (E). Quantification of egg laying (F) and percentage of eggs that failed to hatch 24 hours after being laid (G). All graphs (except C) show means ± SEM. P****≤0.0001 (A-C, one-way ANOVA); (D-G Two-way unpaired t test).

To determine whether the reduced mating activity was a result of either decreased male courting or female receptivity in *dSERT^16^*mutants, we paired the mutants with wild type flies. Pairing *dSERT^16^*mutant females with control males significantly reduces sleep compared to *dSERT^16^* male-female coupling. By contrast, we did not detect a statistically significant reduction in sleep when *dSERT^16^* males were coupled with wild type control females (Figure 4B). These behavioral analyses suggested that *dSERT^16^* males, but not females, exhibit defects in sexual behavior.

To more directly test whether *dSERT* mutants would show defects in sexual activity, we measured copulation rates. We did not observe a significant difference in copulation success in *dSERT^16^* females compared to control females when paired with wildtype males. However, when *dSERT^16^*males were paired with wildtype females none were found to copulate within one hour in marked contrast to the majority of control males (Figure 4C).

In flies, copulation and courtship are regulated in part by distinct pathways. We therefore assayed courtship in the *dSERT* mutant. *dSERT^16^* males exhibited significant defects in courtship behavior including increased latencies to initiate orientation (Figure 4D) and wing song (Figure 4E).

As a further test of male sexual behavior, we measured reproductive output in wildtype females paired for 2 days with *dSERT^16^* males. Wild type females paired with *dSERT^16^*males showed a reduction in egg laying rate (Figure 4F) as well as an increase in the proportion of unfertilized eggs (Figure 4G). These results indicate that while *dSERT^16^* males fail to copulate within one hour, over an increased time-period mating can occur, albeit with reduced fecundity compared to controls.

### Loss of *dSERT* reduces food intake

In addition to alterations in sexual activity, common clinical side effects of SSRIs include changes in food intake, weight gain and gastrointestinal distress (37, 50). Previous work has demonstrated that starvation induces sleep loss in *Drosophila* and increases activity to search for food (51). We therefore tested whether starvation was sufficient to suppress sleep in *dSERT^16^*mutants. Consistent with previous studies, control flies dramatically suppressed their sleep when starved on agar medium compared to their sleep on baseline or recovery days (Figure 5A). By contrast, starvation did not have a significant effect on sleep in the *dSERT^16^* mutants (Figure 5A). Furthermore, we directly tested the feeding in these *dSERT* mutants and found that after 24 hours of starvation food uptake in *dSERT^16^*mutants was significantly decreased compared to controls (Figure 5B). Taken together these results indicate that loss of *dSERT* not only enhances sleep but reduces food intake.

**Figure 5.**
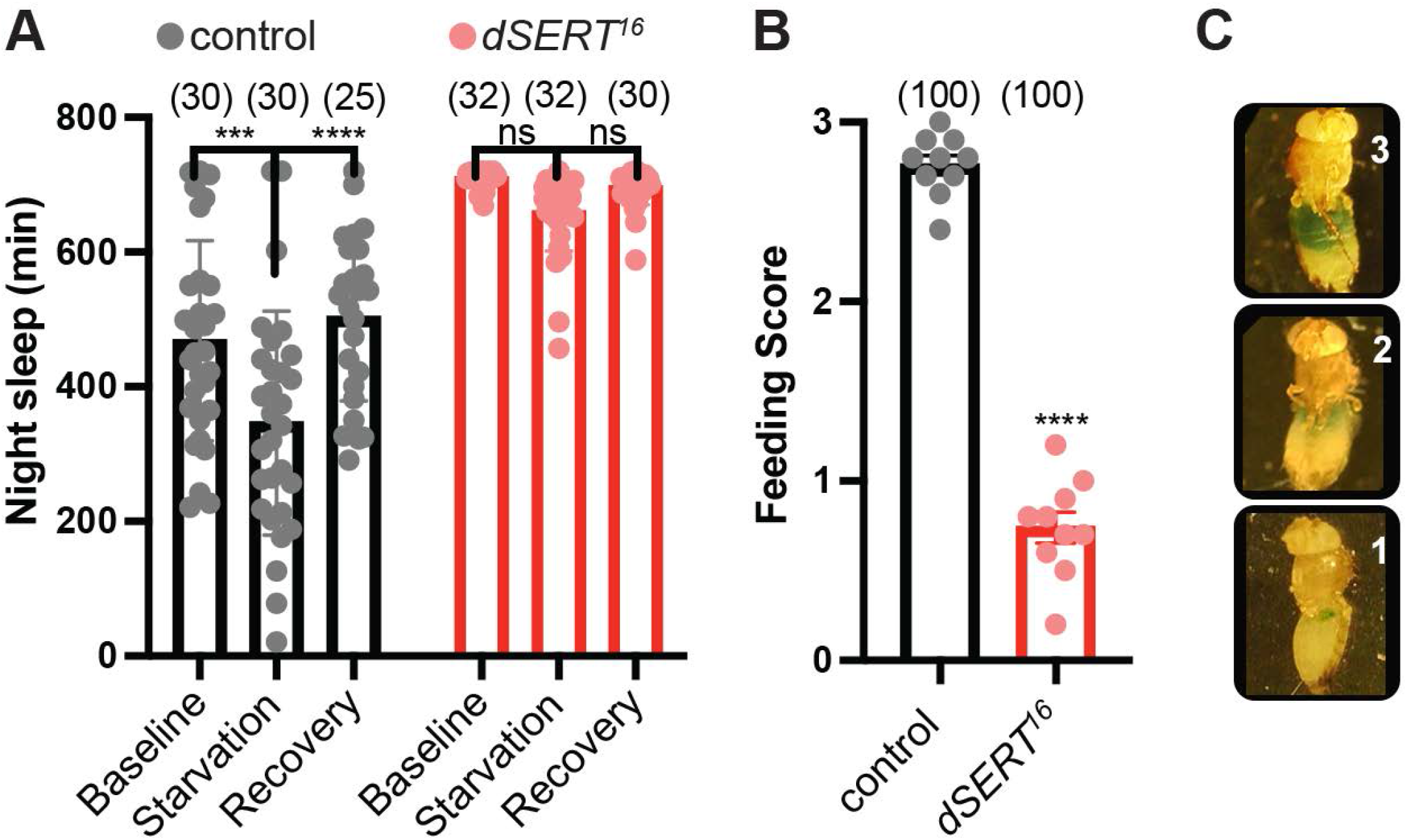
*dSERT^16^* mutants exhibit defects in feeding. (A) Quantification of nighttime sleep for wild-type *w^1118^* (grey) and *dSERT^16^* mutants (red) over a 3-day period that began 1 day after initial loading into DAMs tubes. Day 1 (Baseline): flies kept on standard food. Day 2 (Starvation): flies transferred to agar for food deprivation. Day 3 (Recovery): flies transferred to fresh food. (B) Food uptake is significantly reduced in starved *dSERT^16^*mutants (red) compared to controls (grey). (C) Representative images depict visual feeding score used to assay food uptake. All graphs show individual datapoints and group means ± SEM. P****≤0.0001 (two-way ANOVA (A), two-way unpaired t test (B)).

### Transgenic expression of *dSERT* in serotonergic neurons is sufficient to rescue nighttime or daytime sleep defects

To further confirm that defects seen in *dSERT^16^* mutants are due to a loss of *dSERT* we used “genetic rescue” and expressed a wild type *dSERT* transgene in the *dSERT^16^* background, focusing primarily on defects in sleep. We first used the broad serotonergic driver *TRH-Gal4* (52) to express *UAS-dSERT*. Although, sleep in the fly has been generally approached as a single behavior, some genes and environmental factors can preferentially affect one phase (53–56). Expression of *UAS-dSERT* using *TRH-Gal4* did not fully reverse the increase in total sleep across the 24-LD cycle in *dSERT^16^* mutants (Figure 6A-B). However, separately analyzing day versus night sleep shows that expression of *dSERT* using *TRH-Gal4* is sufficient to restore the daytime sleeping pattern to control levels (Figure 6C). By contrast, night sleep was not altered in the *dSERT^16^* mutants by *TRH-Gal4>UAS-dSERT* (Sup Figure 3A). We further analyzed whether sleep architecture could be rescued. Expression of *UAS-dSERT* by *TRH-Gal4* in *dSERT^16^* mutant was also able to restore bout frequency to the level of control flies for daytime sleep (Figure 6D). Conversely, we did not observe rescue of the nighttime sleep consolidation phenotype (Sup Figure 3B).

**Figure 6.**
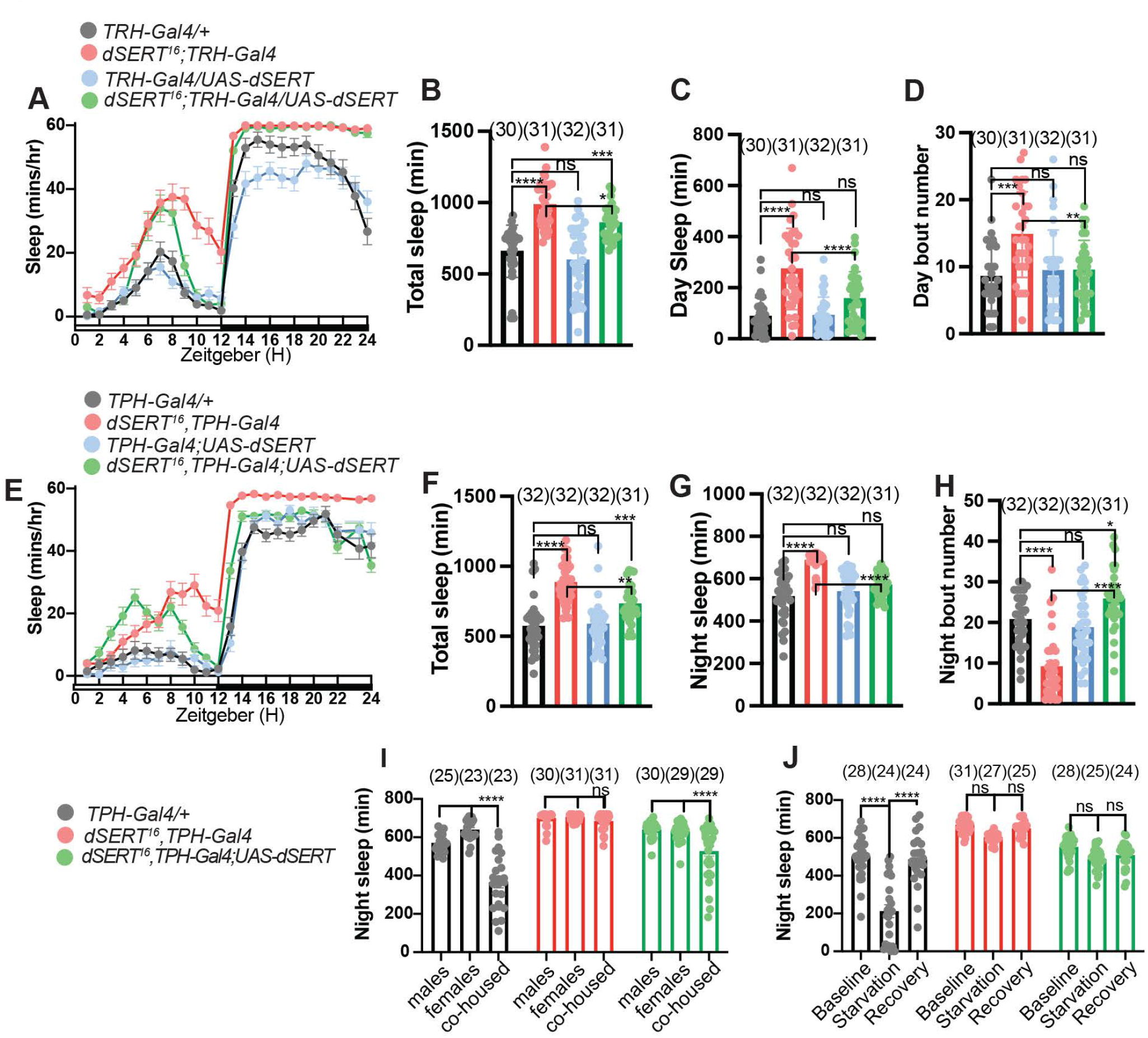
Transgenic expression of *dSERT* with *TRH-Gal4* or *TPH-Gal4* is sufficient to rescue daytime defects or nighttime defects respectively in *dSERT^16^* mutants. (A) Hourly sleep traces in wild-type *w^1118^;TRH-Gal4* (grey), *dSERT^16^; TRH-Gal4* (red), *TRH-Gal4/UAS-dSERT* (blue), and *dSERT^16^; TRH-Gal4/UAS-dSERT* (green) flies. Quantifications of total sleep (B), daytime sleep (C), and daytime bout number (D). (E) Hourly sleep traces in wild-type *w^1118^;TPH-Gal4* (grey), *TPH-Gal4*, *dSERT^16^*(red), *TPH-Gal4;UAS-dSERT* (blue), and *TPH-Gal4*, *dSERT^16^; UAS-dSERT* (green) flies. Quantifications of total sleep (F), nighttime sleep (G) and nighttime bout number (H). (I) Quantification of nighttime sleep for males and females individually housed in DAMs tubes for control wild-type *w^1118^;TPH-Gal4* (grey), *TPH-Gal4*, *dSERT^16^* (red), and *dSERT^16^; TRH-Gal4/UAS-dSERT* (green) flies. Following 2 days of individual housing male and females were paired together in DAMs tubes and nighttime sleep was quantified. (J) Quantification of nighttime sleep for wild-type *w^1118^;TPH-Gal4* (grey), *TPH-Gal4*, *dSERT^16^* (red), and *dSERT^16^; TRH-Gal4/UAS-dSERT* (green) flies over a 3-day period that began 1 day after initial loading into DAMs tubes. Day 1 (Baseline): flies kept on standard food. Day 2 (Starvation): flies transferred to agar for food deprivation. Day 3 (Recovery): flies transferred to fresh food. Sleep traces show mean ± SEM and graphs show individual datapoints and group means ± SEM. P****≤0.0001 (one-way ANOVA (B-D, F-H) and two-way ANOVA (I-J)).

To extend our analysis we tested another broad serotonergic driver, *designated as “TPH-Gal4”* (57). While both *TRH-Gal4* and *TPH-Gal4* include fragments of the *Tryptophan Hydroxylase* gene, they have been reported to exhibit differences in expression (58) including a relatively lower of expression for *TRH-Gal4* in processes that innervate the mushroom bodies for. Similar to *TRH-GAL4*, total sleep levels were not fully rescued using *TPH-Gal4 expressing UAS-dSERT in the dSERT^16^ background* (Figure 6E, F). However, in contrast to *TRH-GAL4*, analysis of nighttime sleep demonstrates that rescue via *TPH-Gal4* is sufficient to restore nighttime sleep levels to that of controls (Figure 6G). Moreover, unlike *TRH-Gal4*, *TPH-Gal4* did not rescue daytime levels (Sup Figure 3D). Further analysis of sleep architecture revealed that expression of *UAS-dSERT* with *TPH-Gal4* in *dSERT^16^* mutants was able to rescue the nighttime consolidation phenotype (Figure 6H). while daytime architecture did not significantly differ from the *dSERT^16^*mutant (Sup Figure 3E).

To further extend our rescue experiments to sexual and feeding in the context of sleep, we repeated our experiments using either starvation or co-housing, focusing on nighttime sleep and rescue with *TPH-Gal4.* Consistent with the results shown in Figure 4, control flies reduced nighttime sleep when male and females were paired and *dSERT^16^* mutants did not show any significant decrease in sleep in co-housing conditions (Figure 6I). However, flies expressing *UAS-dSERT* under the control of the *TPH-Gal4 driver*, significantly decrease their nighttime sleep in co-housing conditions compared to sleep levels of individual male or females (Figure 6I). While we cannot yet clearly parse the specific effects of rescue on sexual behavior versus sleep, these data suggest that expression of *UAS-dSERT* with *TPH-Gal4* restored the relative balance of sexual behavior vs. sleep and most likely rescued both behaviors.

We next tested if genetic rescue using *TPH-Gal4* could also restore sleep to wild type levels under starvation conditions. When starved, nighttime sleep was again decreased in control flies but was not significantly reduced in *dSERT^16^* mutants (Figure 6J). However, under starvation conditions, expression of *dSERT* under the control of *TPH-Gal4* in *dSERT^16^* mutants did not significantly decrease nighttime sleep when starved (Figure 6J). Overall, these results demonstrate that the sleep defects in *dSERT^16^*mutants can genetically rescued, confirming that this phenotype is due to disruption of the *dSERT* gene. In addition, they strongly suggest for the first time that distinct serotonergic circuits may regulate day and night sleep separately. Moreover, they suggest that while the serotonergic circuits that control nighttime sleep and sexual behavior may share at least some of the same pathways, the serotonergic pathways responsible for regulating feeding and nighttime sleep are likely to be independent.

## Discussion

We find that dSERT functions as a critical modulator of sleep, with loss of *dSERT* causing a severe increase in both daytime and nighttime sleep. Given that dSERT acts as the primary mechanism by which serotonin is cleared from the extracellular space, it is likely that *dSERT* mutants have an increase in the amount of serotonin available for neurotransmission. Their behavior is consistent with phenotypes caused by other changes in serotonergic signaling: feeding the serotonin precursor 5-HTP or overexpression of the rate-limiting enzyme *TRH* in dopaminergic and serotonergic cells (16) (*Ddc-Gal4*) increases in total sleep, similar to *dSERT* mutants. Conversely, a decrease in total sleep duration is exhibited by *TRH* mutants (20).

A role for serotonin has also been suggested to contribute to male courtship and mating on based in part on studies of the gene *fruitless* (*fru*) (59, 60). Male specific Fru^M^ positive abdominal-ganglionic neurons that innervate the reproductive organs have been shown to be serotonergic and *fru* mutants demonstrate defects in courting and mating events (59, 60). Furthermore, consistent with our findings, activation of serotonergic neurons causes defects in male copulation behavior (58).

Previous studies have also addressed the complex relationship between increased serotonin signaling and feeding behavior in *Drosophila*. The use of either *TPH-Gal4* or *TRH-Gal4* to thermogenetically depolarize a broad population of serotonergic cells can significantly decrease food intake (58, 61), whereas the activation of a specific subset of serotonergic cells drives a contradictory behavior and induces feeding in sated flies (61).

Our work demonstrates that the increase in overall sleep in *dSERT^16^* mutants is mediated by two different changes in sleep architecture that differ during daytime versus nighttime periods. During the nighttime the increase in sleep in *dSERT^16^* flies is characterized by significantly decreased bout number (consolidated sleep) whereas during daytime sleep bout number is increased. The differential rescue nighttime or daytime sleep via *TPH-Gal4* or *TRH-Gal4* respectively, support the idea that different serotonergic circuits may regulate nighttime vs. daytime sleep. Previous work has shown that knockdown of *TRH* in the dorsal paired median (DPM) neurons that innervate the MB causes a decrease in nighttime sleep (19). We therefore speculate that changes in nighttime sleep caused by dSERT may be mediated primarily through an increase in serotonergic signaling to the mushroom body (MB). Consistent with our findings that *TRH-Gal4* could rescue only daytime sleep, previous work has reported that the expression pattern of the *TRH-Gal4* shows relatively little labeling of processes that innervate the MBs (18). We have confirmed that *TRH-Gal4* shows less labeling to the of the MB compared to *TPH-Gal4* (Sup Figure 3C, F). Further evidence for the possibility that the MBs mediate nighttime sleep include the observation the *d5-HT1A* receptor mutant exhibits a decrease in nighttime sleep, nighttime bout length, and an increase in nighttime bout number, and that all of these behaviors can be genetically rescued via expression of a *d5-HT1A* transgene in the MBs (16). Previous work has demonstrated the MBs are dispensable for sleep-feeding interactions (51), consistent with our results showing that even though transgenic expression of *dSERT* with *TPH-Gal4* could restore nighttime sleep levels, it did not rescue the a decrease in sleep via starvation seen in wt flies. Together these data suggest that the increased and consolidated nighttime sleep seen in *dSERT^16^* mutants is the result of an increase in signaling through the d5-HT1A receptor caused by an increase in extracellular serotonin in the MBs.

We speculate that some aspects of the daytime dSERT sleep phenotype, such as the increased frequency of daytime sleep bouts may be mediated by the ellipsoid body and the d5-HT7 receptor. Activation of EB neurons leads to an increase in the number of sleep episodes specifically during the daytime (18). *d5-HT7* mutants and treatment with a 5-HT7 antagonist similarly causes a reduction in sleep episode frequency (18). However, it is likely that the daytime *dSERT* sleep phenotype is dependent on other circuits and/receptors, since *dSERT* exhibit an increase in sleep amount that was not detected by disruption of *d5-HT7* (18). We plan to use the *dSERT* mutants to further map serotonergic circuits that may differentially daytime vs nighttime sleep.

Studies in mice underscore the complexity of serotonin’s effects on sleep. For example, an increase in serotonergic signaling caused by increased activity of raphe serotonergic cells reduces REM sleep (62)(63)(64) while an increase in serotonin levels caused by a *SERT KO* mouse has the opposite effects and increases REM (65, 66). Knockouts of either 5-HT1B or 5-HT1A also exhibit more REM (67, 68), but the increase in REM caused by the SERT KO can be reduced by developmental blockade of 5-HT1A (66).

In humans, the complex relationship between sleep, depression, and the effects of SSRIs underscores the importance of understanding the role of SERT in sleep. Variation in the 5’ flanking region of the human *SERT* gene have been associated with a change in SERT expression levels *in vitro* (69), that is linked to the efficacy of SSRIs (70) and sleep deprivation (71). In addition to sleep, SSRIs impact other critical behaviors such as feeding and mating. SSRIs can induce hypophagic effects and interfere with normal feeding behavior (72–75), and sexual dysfunction is a well-documented side effect of SSRI treatment (35, 36). Our work begins to address the competition between the pathways that regulate these effects in the context of *Drosophila* sleep, as we demonstrated that loss of *dSERT* induces an increased sleep drive that outweighs the propensity for both normal feeding and mating behaviors (Figure 5). We speculate that further studies of *dSERT* mutants may help to uncover the relationship between other serotonergic pathways that regulate other complex behaviors common to mammals and flies such as aggression and may be used further as a genetic model for the behavioral effects of SSRIs (52,76–80).

## Materials & Methods

### Fly Strains

Fly stocks were reared on standard cornmeal media (per 1L H20: 12 g agar, 29 g Red Star yeast, 71 g cornmeal, 92 g molasses, 16mL methyl paraben 10% in EtOH, 10mL propionic acid 50% in H20) at 25 C with 60% relative humidity and entrained to a daily 12 h light,12 h dark schedule (12hr:12hr LD). *w^1118^* flies were obtained from Dr. Seymour Benzer. *UAS-dSERT* (BL24464) and *UAS-MCD8::GFP* (BL32185) were obtained from the Bloomington *Drosophila* Stock Center. *TPH-Gal4* was from Dr. Jongkyeong Chung(57).

### P-element mutagenesis

The *dSERT* alleles were generated by imprecise excision of the *P{XP}d04388* transposon that is inserted 514 bases upstream of the *dSERT* gene on the second chromosome. First, the *P{XP}d04388* line was backcrossed for five generation to the *w^1118^* line of the Scholz lab to isogenize the genetic background. Second, females of the *P{XP}d04388* were crossed to *w^1118^*; *CYO/+; [Δ2-3; Ki]/TM2* males. Next, *w^1118^; P{XP}d04388/CYO*; [Δ 2-3; Ki]/+ males were mated with *P{XP}d04388* females. In the next generation, 200 single male flies carrying possible deletion of the *P{XP}d04388* over *P{XP}d04388* were crossed to *Df(2R) PX2/CYO* females that carry a genomic lesion including the complete *dSERT* gene. The new dSERT alleles complemented the lethality associated with the deficiency. The males of the F1 generation were screened for loss of the P-element insertion, e.g. white-eyed male flies carrying the *CYO* allele. 89 lines were established as stocks. The new alleles were analyzed for genomic lesions using PCR. The following primers were used to confirm lesion and amplify DNA for sequencing: 5’-TGACCCACTAAATGCCATGA-3’ and 5’ CCAGAAAAAGCGAAATCTGC-3’.

### Quantitative Real Time PCR (qRT-PCR)

Total RNA of 30 flies was isolated using Trizol™ method followed by a DNAse digest for 30 min at 37° C. To synthesize cDNA, the SuperScript^TM^ II Reverse Transcriptase (Invitrogen^TM^) and oligo^dT^ primers were used according manufacturer instruction. For qRT-PCR analysis the MESA BLUE qPCR MasterMix Plus for SYBR® Assay (Eurogentec) with 100 ng cDNA as template was used. The detection was performed using the iCycler iQ5 Multicolour Real-Time PCR Detection System. The data were analyzed using the ΔΔCt method(81). The NormFinder software(82) was used for control primers selection. The following control primers recognizing *RpLPO* were selected: 5‘-CAG CGT GGA AGG CTC AGT A-3‘ and 5’-CAG GCT GGT ACG GAT GTT CT-3’. For *dSERT* the following primers recognizing sequences in the third and fourth exon were used: 5’-GTTGCCTCAGCATCTGGAAG-3‘ and 5’-CAGCCGATAATCGTGTTGTA-3’. Data are shown as fold changes in *dSERT* relative to the controls.

### Western blot analysis

To isolate proteins of fly heads, 500-1000 flies were frozen in liquid nitrogen and heads were separated with a sieve. The heads were pulverized with a sterile pestle and resuspended in buffer containing 10mM NaCL, 25 mM Hepes, pH 7.5, 2mM EDTA and 1x cOmplete^™^ Mini protease inhibitors (Merck). The suspension was kept on ice for 10 min with gentle mixing before spinning it at 18300 x g at 4°C for 15 min. The remaining pellet was resuspended in a buffer containing 10mM NaCL, 25 mM Hepes, pH 7.5, 1x cOmplete^™^ Mini protease inhibitors (Merck) and 0.5% CHAPS and incubated at 4°C for 10 minutes with gentle mixing. The solution was centrifuged at 3000 x g for 5 min to recover membrane extract. 20 μg protein of each sample were used for western blotting according standard procedures. The membrane was blocked in 5% milk and the primary rabbit anti-dSERT antibody (Giang et al., 2011) were used at a dilution of 1: 20 000 and mouse anti-β-Actin (mAbcam 8224) at 1:10 000.

### Behavioral Analysis

#### Sleep

Sleep was measured as previously described(83). In brief, 3-5 day old female flies (co-housing experiments also utilized male flies) were individually loaded into 65mm long glass tubes and inserted into *Drosophila* activity monitors to measure locomotor activity (Trikinetics Inc; Waltham MA, USA) and periods of inactivity for at least 5 minutes were classified as sleep. Custom Visual Basic scripts(83) in Microsoft Excel were used to analyze Trikinetics activity records for sleep behaviors. MB5 monitors (Trikinetics Inc; Waltham MA, USA) were used for multibeam monitoring experiments.

#### Starvation

For starvation and sleep experiments 3-5 day old, mated females were loaded into DAMs tubes with standard food for acclimation. After 1 day of acclimation, baseline sleep was measured for 24 hours (Day 1). On the experimental day (Day 2) flies were transferred to tubes containing 1% agar at ZT7 to be starved for 17 hours. Flies were then transferred back to fresh tubes containing standard food at ZT0 (Day 3) and activity was recorded for 24-hour recovery period.

#### Co-housing

For co-housing sleep experiments males and females were loaded individually into DAMs tubes and baseline sleep was recorded for 2 days. On experimental day at ZT0 flies were combined into male-female pairs (either same genotype or mixed as specified in the text) in new DAMS tubes and sleep was recorded for 24 hours. The following day at ZT0 flies were separated back into individual tubes and activity was recorded for 24-hour recovery period.

#### Grooming

For grooming experiments, 3–5-day old, mated females were observed in individual chambers using a Dinolite USB video camera AM7023CT. A single grooming event was scored when a fly rubbed its legs over its head or abdomen or rubbed legs together. For each fly, grooming was analyzed over three separate 2 min periods and quantified as an average number of grooming events per minute. Each scoring period was started by the observation of an initial single grooming event.

#### Negative Geotaxis

For negative geotaxis experiments, 3 groups of 10 flies were placed in two empty 28.5 x 95 mm polystyrene vials that are vertically joined by tape facing each other. After gently tapping the vials, the flies were allowed to climb for 10s. Each group was scored for the number of flies that climbed above the 8cm mark compared to those remaining below the 8cm mark. Each group was tested 5 times (with one minute recovery between each trial), and the average for those five trials was quantified.

#### Locomotor Activity and Circadian Rhythmicity

Experiments were performed with 3-5-day-old females in *Drosophila* activity monitors (Trikinetics Inc; Waltham MA, USA) that were first entrained for 2-3 days in 12hr:12hr LD. For DD analysis, data were analyzed for 3-6 days from the second day in DD. Data analysis was done with Faas software 1.1 (Brandeis Rhythm Package, Michel Boudinot). Rhythmic flies were defined by χ^2^ periodogram analysis with the following criteria: power ≥20, width ≥2h with selection of 24hr ± 8hr upon period value. Power and width are the height and width of the periodogram peak, respectively, and give the significance of the calculated period. Mean daily activity (number of events per 0.5hr ± SEM) is calculated over 3-4-day period in LD or DD conditions. Morning anticipation index was calculated as(84) (activity for 3hrs before lights-on)/(activity for 6hrs before lights-on) and evening anticipation index as (activity for 3hrs before lights-off)/(activity for 6hrs before lights-off).

#### Courtship & Copulation Assays

Virgin females were aged 3-5 days in groups ∼10 and naïve males 4 days individually. Single pairs of female and male flies were gently aspirated into a chamber of a mating wheel and video recording was started. The percent of males achieving copulation within 1 hour was measured as well as the time elapsed between the introduction into the chamber and the male display of courtship steps such as orientation and wing extension.

#### Egg Laying and Hatchability

Five-day old virgin females fed with wet yeast for one day were placed with males (5 females to 10 males) in one bottle for egg laying on molasses plates over 2 days (with removal and replacement of plates every 24 hours). Once molasses plates were removed, they were kept in sealed Tupperware and after 24 hours in 25°C the number of unhatched eggs were manually counted under a microscope.

#### Feeding Assays

Groups of 10 flies were starved for 24 hours on 1% agar medium as a water source. After starvation, flies were transferred to blue food (I will add food recipe) and allowed to feed for 15 mins. The amount of blue food dye ingested was measured using visual inspecting feeding scores as previously described(61). From 0 being no dye visible in the abdomen to 3, the abdomen is swollen and filled with dye. The average feeding score was quantified by averaging the feeding scores of each fly in the vial.

### Immunohistochemistry

Immunostaining was performed following a standard procedure comprising brain dissection, fixation in 3% glyoxal for 25 min, blocking in PBTG (PBS plus 0.2% Triton, 0.5% BSA and 2% normal goat serum), and primary and secondary antibody staining diluted in PBTG. The following primary antibodies were used: rabbit anti-GFP (1:500, Invitrogen) and mouse anti-DLG (1:20, 4F3 -Developmental Study Hybridoma Bank). Alexa Fluor 488 and Alexa Flour 555 goat secondary antibody (1:500; Invitrogen) were used as secondary antibodies. Confocal images were obtained using a Zeiss LSM 880 confocal microscope with Zen software. Images were processed using Fiji/Image J software.

### Statistics

Data were analyzed in Prism 9 (GraphPad; San Diego CA, USA). Group means were compared using two-tailed t tests or one-or two-way ANOVAs, with repeated-measures where appropriate, followed by planned pairwise comparisons with Holm-Sidak multiple comparisons tests. Sample sizes for each experiment are depicted in each figure panel or in the appropriate figure legend. All group averages shown in data panels depict mean ± SEM.

## Acknowledgements

We thank Drs Jongkyeong Chung for sharing fly lines. We thank Dr. Ceazar Nave and Prabhjit Singh in J.D.’s laboratory for technical support and discussion. We thank Dr. Tim Lebestky and Mikhayla Armstrong for their early contributions to the analysis of the *dSERT* mutants.

---

**Supplemental Figure 1.**
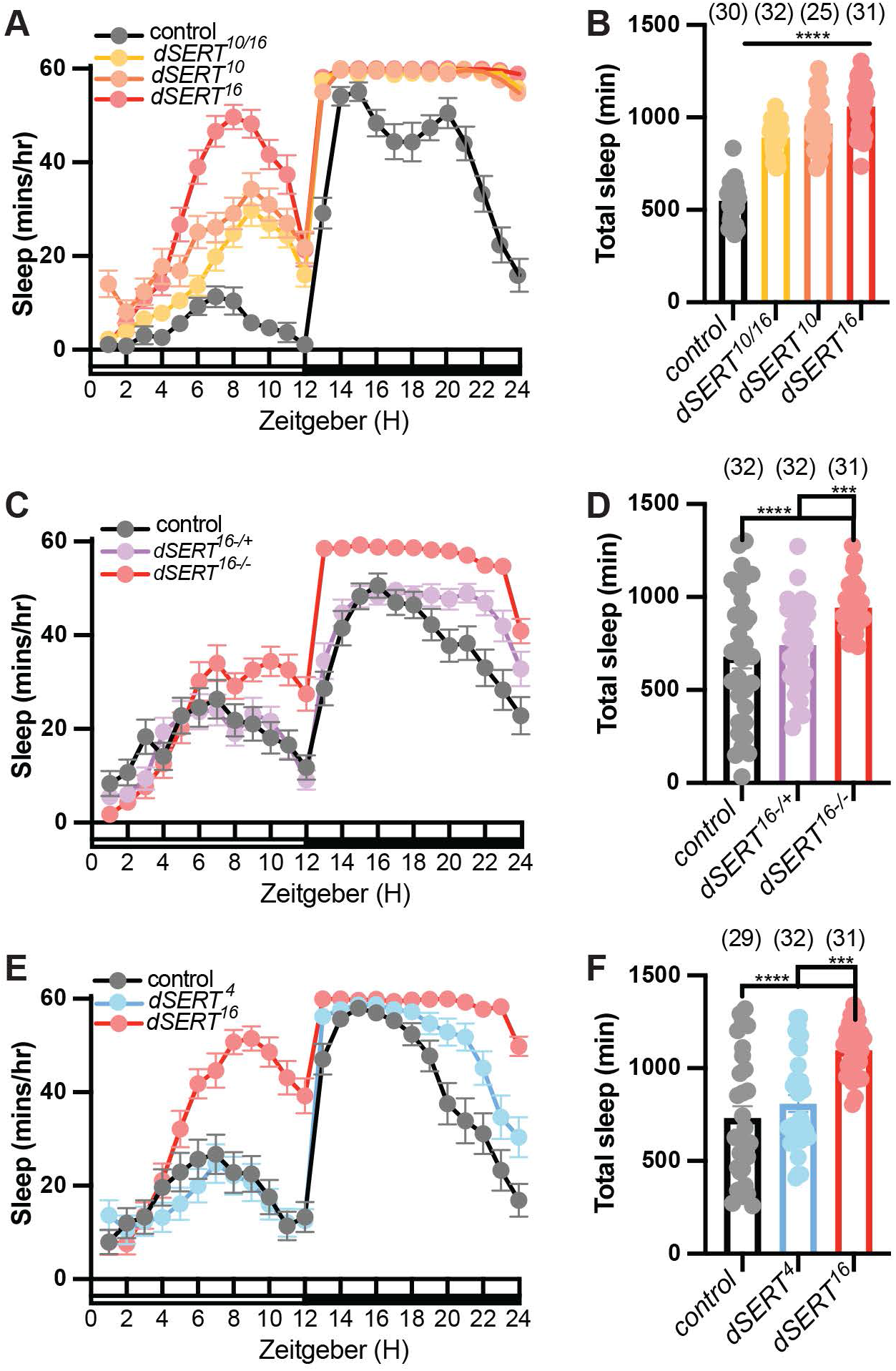
Genetic controls. Hourly sleep traces (A) and quantification of total sleep (B) in control *w^1118^* (grey), transheterozygous *dSERT^10^/dSERT^16^* mutants (yellow), *dSERT^10^* mutants (orange), and *dSERT^16^* mutants (red). Hourly sleep traces (C) and quantification of total sleep (D) in control *w^1118^* (grey), *dSERT^16^*heterozygotes (purple), and *dSERT^16^* homozygous mutants (red). Hourly sleep traces (E) and quantification of total sleep (F) in control *w^1118^* (grey), *dSERT^4^* revertants (blue), and *dSERT^16^* mutants (red). Sleep traces show mean ± SEM and graphs show individual datapoints and group means ± SEM, p≤0.0001****, one-way ANOVA, Holm-Sidak multiple comparisons tests.

**Supplemental Figure 2.**
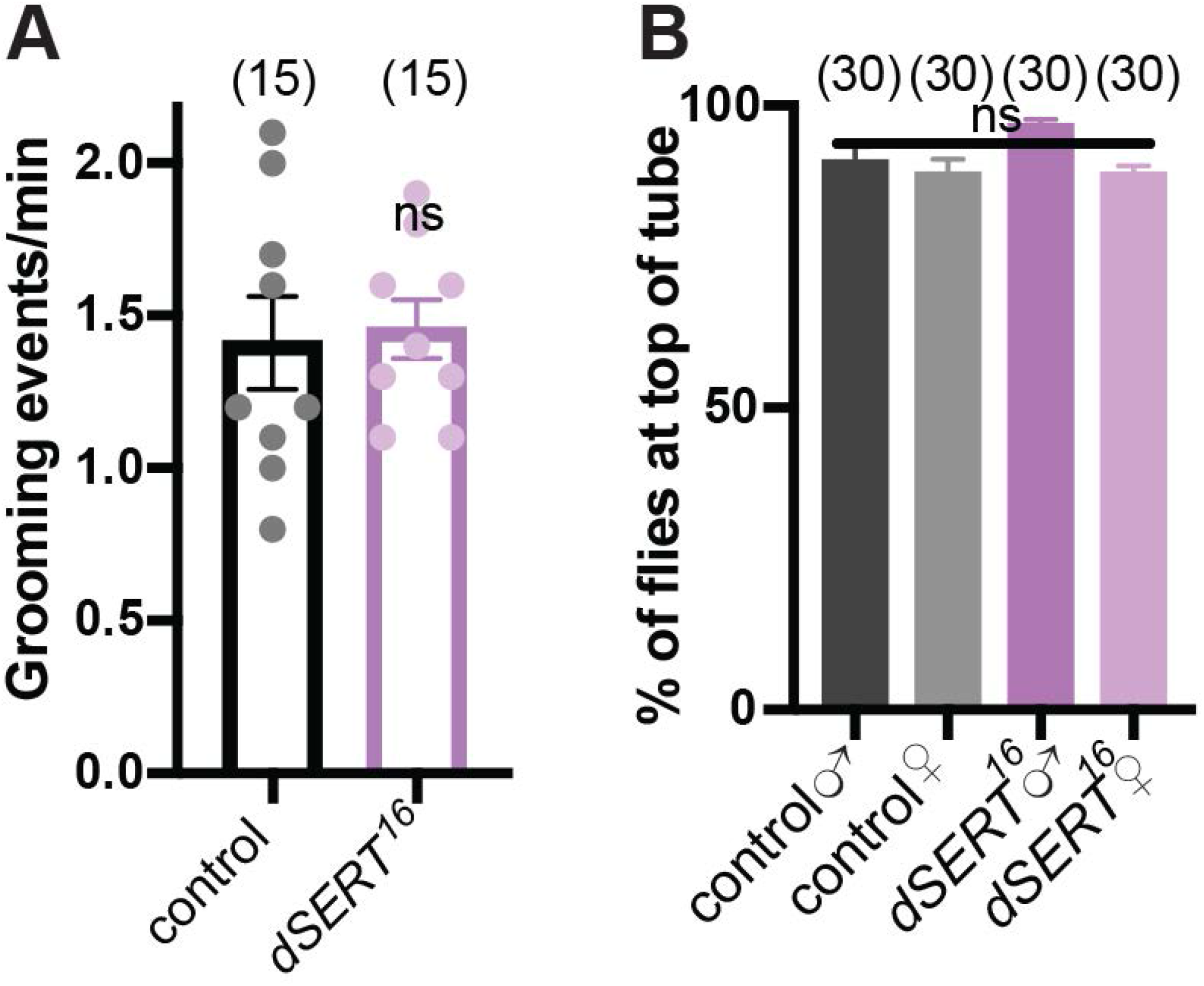
d*S*ERT16 mutant sleep phenotype is not an artifact of additional amine-linked behaviors. (A) *dSERT^16^* mutants (purple) show no change in grooming behavior compared to wild-type *w^1118^* flies (grey). Mean± SEM, unpaired Student’s t-test. (B) Male and female *dSERT^16^* mutants behave indistinguishably from control flies in negative geotaxis assays. Mean ± SEM, one-way ANOVA.

**Supplemental Figure 3.**
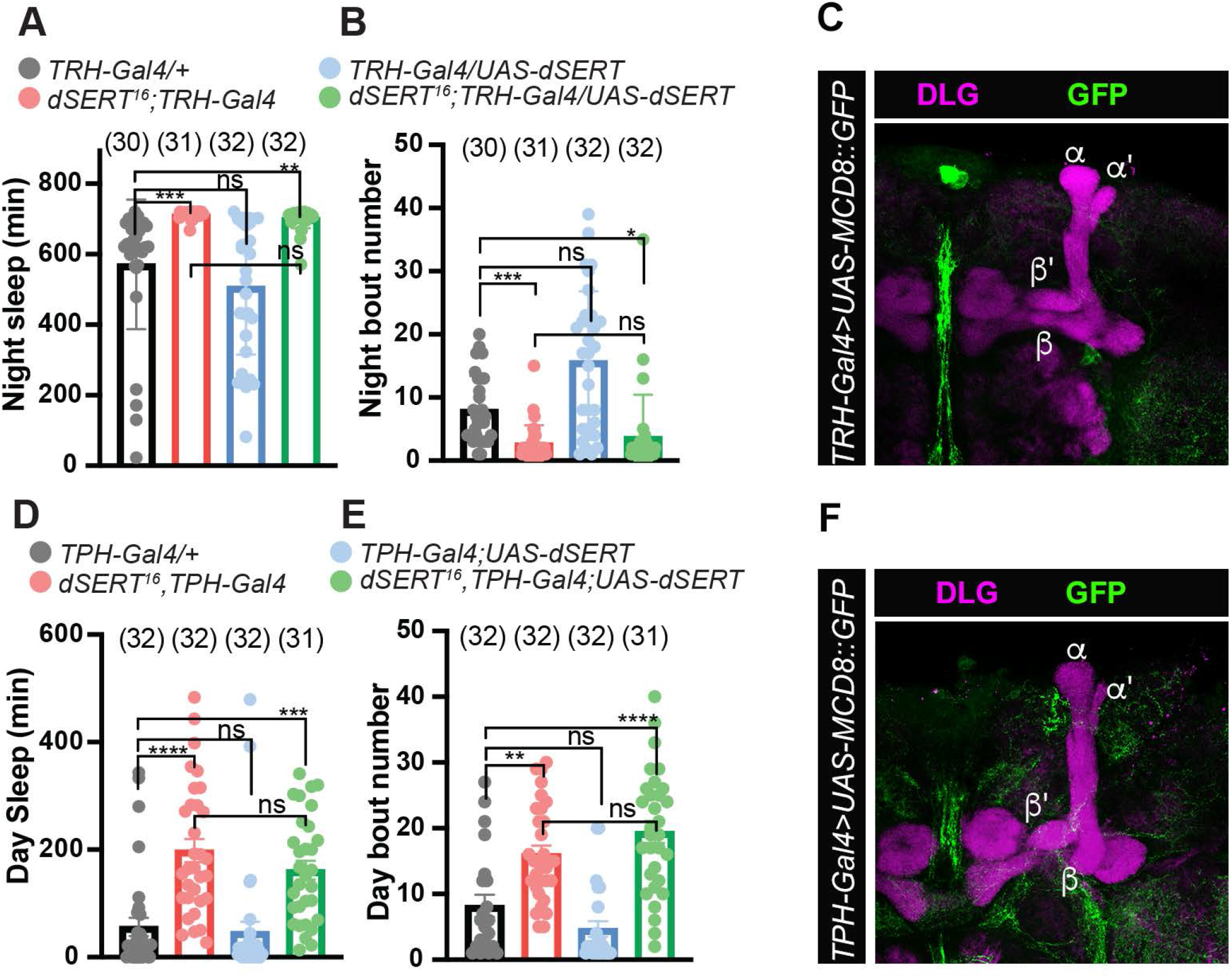
Quantification of nighttime sleep (A) and nighttime bout frequency (B) in wild-type *w^1118^;TRH-Gal4* (grey), *dSERT^16^; TRH-Gal4* (red), *TRH-Gal4/UAS-dSERT* (blue), and *dSERT^16^; TRH-Gal4/UAS-dSERT* (green) flies. (C,F) Representative pictures show expression of UAS-MCD8::GFP (green) driven by *TRH-Gal4* (C) or *TPH-Gal4* (F) and DLG (magenta) in mushroom bodies. The different lobes of the mushroom body are labeled in white. Quantification of daytime sleep (D) and daytime bout frequency (E) in wild-type *w^1118^;TPH-Gal4* (grey), *TPH-Gal4*, *dSERT^16^* (red), *TPH-Gal4;UAS-dSERT* (blue), and *TPH-Gal4*, *dSERT^16^; UAS-dSERT* (green) flies. All graphs show individual datapoints and group means ± SEM. P****≤0.0001 (one-way ANOVA).

